# Subject-independent decoding of affective states using functional near-infrared spectroscopy

**DOI:** 10.1101/228007

**Authors:** Lucas R. Trambaiolli, Juliana Tossato, André M. Cravo, Claudinei E. Biazoli, João R. Sato

## Abstract

Affective decoding is the inference of human emotional states using brain signal measurements. This approach is crucial to develop new therapeutic approaches for psychiatric rehabilitation, such as affective neurofeedback protocols. To reduce the training duration and optimize the clinical outputs, an ideal clinical neurofeedback could be trained using data from an independent group of volunteers before being used by new patients. Here, we investigated if this subject-independent design of affective decoding can be achieved using functional near-infrared spectroscopy (fNIRS) signals from frontal and occipital areas. For this purpose, a linear discriminant analysis classifier was first trained in a dataset (49 participants, 24.65±3.23 years) and then tested in a completely independent one (20 participants, 24.00±3.92 years). Significant balanced accuracies between classes were found for positive vs. negative (64.50 ± 12.03%, p<0.01) and negative vs. neutral (68.25 ± 12.97%, p<0.01) affective states discrimination during a reactive block consisting in viewing affective-loaded images. For an active block, in which volunteers were instructed to recollect personal affective experiences, significant accuracy was found for positive vs. neutral affect classification (71.25 ± 18.02%, p<0.01). In this last case, only three fNIRS channels were enough to discriminate between neutral and positive affective states. Although more research is needed, for example focusing on better combinations of features and classifiers, our results highlight fNIRS as a possible technique for subject-independent affective decoding, reaching significant classification accuracies of emotional states using only a few but biologically relevant features.

## 1. Introduction

Multivariate brain decoding (MBD) allows the inference of mental states based on diverse features of brain signals (1). In comparison to conventional analytic methods, this approach has the advantage of considering the various brain regions simultaneously, thus providing information on the neural networks of interest as a whole (2). One particular application of MBD is to identify emotional or affective experiences to develop brain-computer interfaces or neurofeedback protocols (3). The clinical application of affective neurofeedback is particularly interesting since it can enable patients to volitionally control abnormal neural activities associated with psychiatric disorders, leading to brain connectivity reorganization and the relief of symptom severity (4, 5).

Conventional clinical neurofeedback approaches are based on subject-specific designs (4). In other words, the patient is first submitted to an initial screening to create a calibration database for the neurofeedback algorithm and, later, starts the proper neurofeedback training. However, although reliable to the participant’s neural data, this initial step leads to long sessions that might increase physical and mental exhaustion and, consequently, reduce the therapeutic benefits (6). Thus, ideally, therapeutic affective neurofeedback should be trained with the first set of volunteers and applied with little or no calibration to new individuals. However, the subject-independent identification of mental states remains an open technical challenge (1).

Subject-independent emotion classification has been previously reported on facial and body expressions (7, 8), and voice tone (7). A critical limitation of those studies is that volunteers can easily handle the records used in order to achieve the desired result, even without changing their affective states. Moreover, such exclusively behavioral measures do not provide neural information, which is crucial to a properly designed neurofeedback system. Thus, investigations using neurophysiological records such as electrophysiology (EEG) and functional magnetic resonance imaging (fMRI) should foster the development of affective interfaces. Using EEG data, for example, different studies using a subject-independent design combined several features and classifiers to predict the affect experienced by new participants, achieving accuracies between 70 and 95% (9–12). Using fMRI data, accuracy varied from 60% to 80% depending on the number of voxels (from 2000 to 4000) included as predictors (13).

Another challenge for fostering clinical affective neurofeedback applications is the high-cost and limited mobility of the so far most accurate imaging measurements (such as those acquired using fMRI) (6). In this context, functional Near-Infrared Spectroscopy (fNIRS) emerges as an attractive neuroimaging technique for affective decoding. This method is based on low-energy light detectors and transmitters for measuring the light absorption through the cortical surface (14). It makes possible to investigate local changes in the concentrations of oxy and deoxyhemoglobin in response to functional brain activity, similarly to the widely applied fMRI BOLD effect (15). Furthermore, fNIRS presents a good trade-off between spatial and temporal resolutions, with low susceptibility to instrumental and biological noise, and considerably lower cost and higher portability when compared to other non-invasive neuroimaging approaches (16). Affective neuroscience experiments are benefited from fNIRS usage due to its reliability to record the prefrontal cortex activation, as well as allowing both restricted and naturalistic emotion-related experiments (16–18). With this in mind, several studies showed a significant subject-specific offline prediction of affective states using different fNIRS protocols (19–23).

Using a fronto-occipital fNIRS setup, our research group achieved 80 to 95% of within-subject affective classification accuracy (22), and developed a functional subject-specific neurofeedback protocol (24). Here, considering the above-mentioned technical challenges for subject-independent affective decoding, we aimed to evaluate whether these fronto-occipital fNIRS signals provide enough information to the subject-independent offline classification of affective states. To achieve this goal, we collected two independent datasets with reactive and active affective tasks. The reactive task was based on the visualization of a set of images from the International Affective Pictures System (IAPS) that were thought to induce positive, negative or neutral affective states (25). Critically, those images were selected in order to balance the arousal dimension for the different valences. In the active task, participants were instructed to imagine personal situations with positive, negative or neutral affective contexts. We expected decoding performances higher than the chance level when comparing two different affective states in each task. Also, based on the affective-workspace model (26, 27), we expected the relevant information used in classification to come from nodes of neural networks mainly comprising frontal and occipital areas (28, 29).

## 2. Methods

### 2.1. Participants

Forty-nine healthy participants (25 female, 24.65±3.23 years) were enrolled in the first part of the experiment. Inclusion criteria were no previous (self-reported) diagnosis of neurological (ICD-10: G00-G99) and/or psychiatric disorders (ICD-10: F00-F99), and normal or corrected-to-normal vision. All participants were attending college or graduated and provided written informed consent to participate in this study.

Two years later, a second (and independent) sample of twenty healthy participants (10 female, 24.00±3.92 years) was collected by a different researcher, and in a separate laboratory (at the same university). The inclusion criteria and the experimental protocol were equal to the first sample. For both cases, ethical approval was obtained from the local Ethics Committee, and no payment was provided to the participants, according to the national rules.

### 2.2. Functional NIRS acquisition

fNIRS measurements were conducted using the NIRScout System (NIRx Medical Technologies, LLC. Los Angeles, California) using an array of optodes (12 light sources and 12 detectors) covering the prefrontal, temporal and occipital areas. Optodes were arranged in an elastic band, with nine sources and nine detectors positioned over the frontal and temporal regions, and three sources and three detectors over the occipital region. Four positions of the International 10-20 System were adopted as reference points during the setup: sensors 1 and 9 were positioned approximately over the T7 and T8 locations, respectively, while the Fpz and Oz positions were in the center of channels 5-5 and 11-11, respectively, as shown in Figure 1. The source-receptor distance was 30 mm for adjacent optodes, and the used wavelengths were 760 and 850 nm. Signals obtained from the thirty-two channels were measured with a sampling frequency of 5.2083 Hz (determined by the maximum sampling frequency of the equipment - 62.5 Hz - divided by the number of sources - 12) using the NIRStar 14.0 software (NIRx Medical Technologies, LLC. Los Angeles, California).

**Figure 1.**
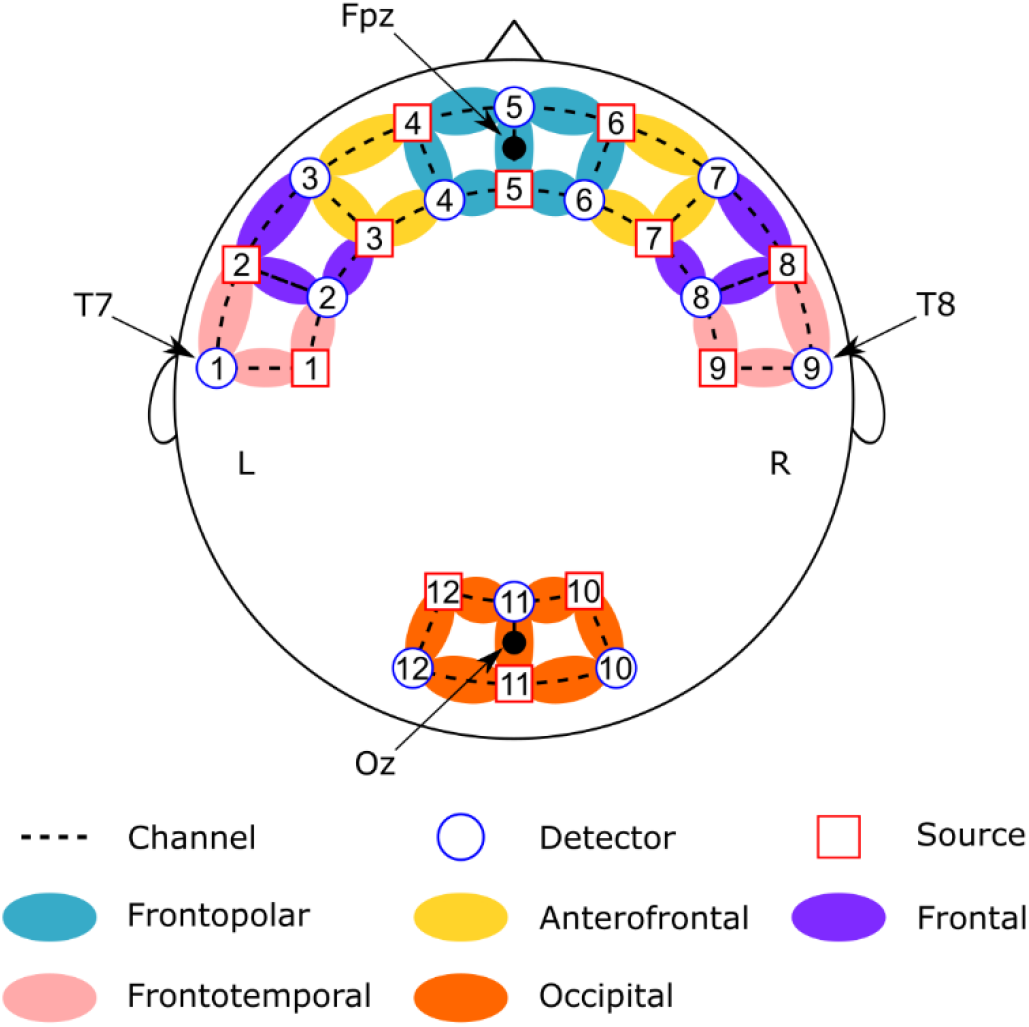
Channel configuration. Red circles represent sources; blue circles represent the detectors and dotted lines the channels. Colors show channel positions according to the International 10-20 system.

### 2.3. Experimental protocol

During the test, participants sat in a padded chair with armrest, positioned 1-meter distance in front of a monitor. They were asked to remain relaxed, with hands within sight resting on the armrests or the table. They were also requested to avoid eye movements, as well as any body movement. The recording room remained dark during registration and subject used earplugs.

#### 2.3.1. Reactive task

For the reactive task, we used images available on the international affective picture system (IAPS) catalog (25). First, images were filtered according to their average values of arousal and then were ranked according to their valence values. We selected the 30 pictures with higher average values of valence, the 30 with lowest values and 60 of intermediate values, as follows:

- Positive pictures (Valence = 7.884±0.220; Arousal = 5.036±0.448): 1811, 2057, 2080, 2209, 5210, 5830, 7200, 2040, 2058, 2091, 2340, 5700, 5833, 7330, 1440, 2045, 2070, 2150, 2347, 5825, 5910, 7502, 1710, 2050, 2071, 2165, 2550, 5829, 5982, 8420;
- Negative pictures (V = 2.007±0.183; A = 5.549±0.339): 2375.1, 3101, 3261, 9181, 9322, 9560, 2703, 3180, 3301, 9185, 9326, 9571, 2095, 2800, 3191, 3350, 9253, 9332, 2205, 3016, 3225, 9040, 9300, 9421, 2345.1, 3062, 3230, 9140, 9301, 9433;
- Neutral pictures (V = 5.234±0.060; A = 3.770±0.813): 2122, 2514, 5520, 7019, 7182, 7550, 2191, 2635, 5531, 7021, 7207, 7632, 2211, 2702, 5532, 7043, 7242, 7830, 1122, 2308, 2745.1, 5533, 7052, 7248, 8065, 1350, 2377, 2850, 5740, 7053, 7249, 1616, 2381, 2870, 5920, 7058, 7365, 1675, 2385, 2880, 6910, 7062, 7497, 1820, 2487, 5395, 7001, 7080, 7500, 1908, 2495, 5471, 7014, 7090, 7506, 2102, 2499, 5510, 7017, 7100.

This task consisted of twenty trials (5 for positive stimuli, 10 for neutral and 5 for negative). For the first 2 seconds of each trial, a white cross was presented in the center of a blank screen. During the next 30 seconds, a new figure was displayed every 5 seconds, totaling six randomly selected figures per trial corresponding to the desired affective class (Figure 2). Presenting a group of images with the same valence ensures the maintenance of cognitive engagement (30) and the required duration to achieve the peak of oxygen concentration change relative to baseline (31). At the end of the trial, a new screen was presented asking the participant to assign a score from 1 to 9 for the subjective valence (1 – extremely negative valence; 9 – highly positive valence) and subjective arousal (1 – lower arousal; 9 – higher arousal) experiences. After this, a blank screen appears for a random duration between 2-4 seconds and participants were instructed to blink and/or move in this period but not in the other phases.

**Figure 2.**
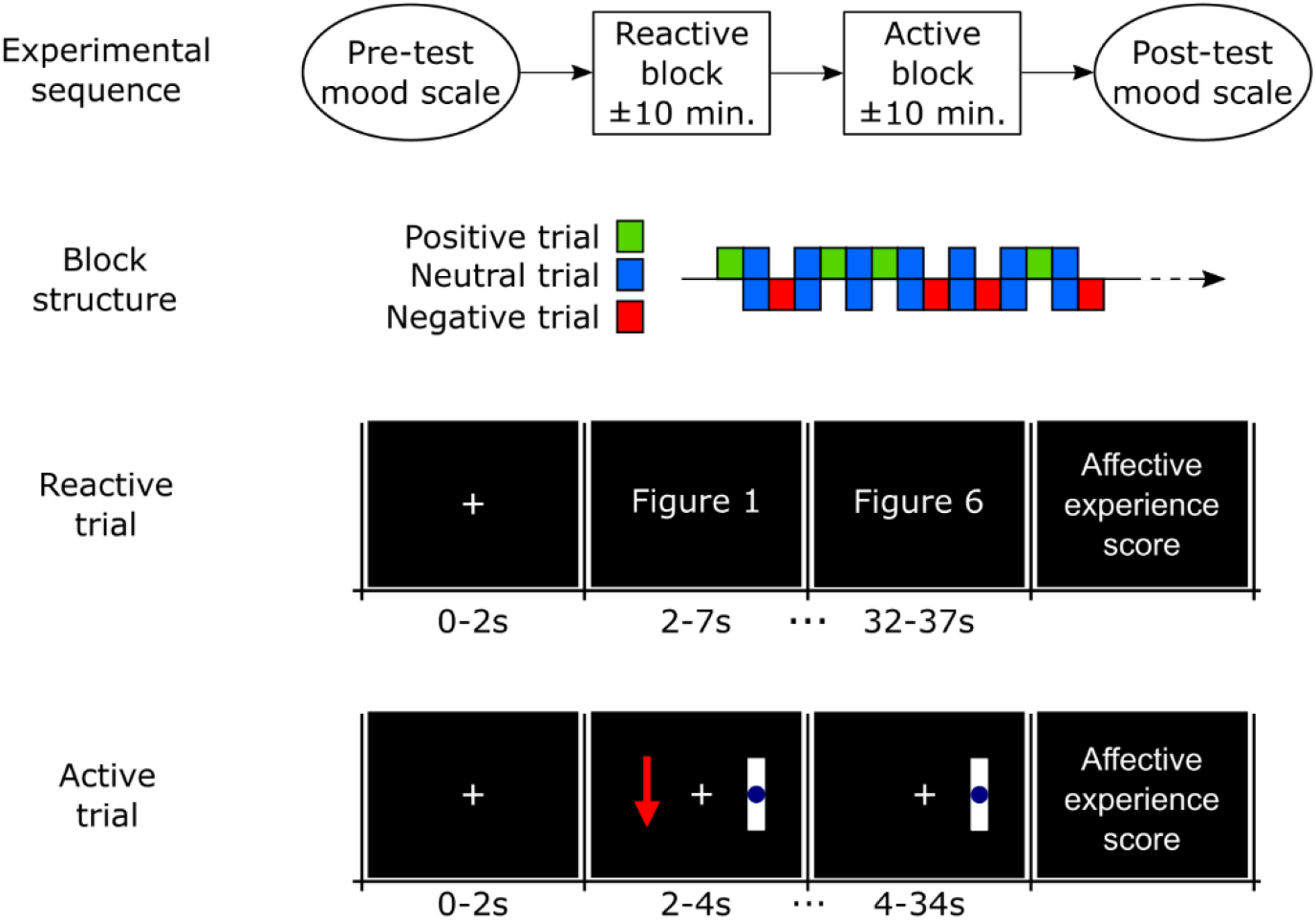
Visual stimuli order in reactive and active trials. The order of trials into the block is random, but always alternating neutral trials with positive and negative affect trials.

#### 2.3.2. Active task

The active task (affective imagination) consisted of twenty trials (5 trials for positive affect, 5 for negative affect and 10 for resting with eyes open, also referred as neutral affect). Each trial started with a baseline period of a blank screen with a white cross in the center. After 2 seconds, the instruction (representing the desired emotion) appeared to the left of the display, remaining on the screen for 2 s (Figure 2). The instruction consisted of either a green arrow pointing up (positive affect), a red arrow pointing downward (negative affect) or a blue circle (neutral affect). For 30 seconds after the instruction was presented, the screen remained unchanged, corresponding to the participant’s affective imagination period. At the end of the trial, a new screen was presented asking the participant to assign a score from 1 to 9 for the subjective valence (1 – extremely negative valence; 9 – highly positive valence) and subjective arousal (1 – lower arousal; 9 – higher arousal) experiences. After this, a blank screen appears for a random duration between 2 to 4 seconds and participants were instructed to blink and/or move in this period.

### 2.4. Data analysis

#### 2.4.1. Preprocessing

Preprocessing was performed using Matlab (Mathworks, MA, USA) with the nirsLAB v2014.12 toolbox (NIRx Medical Technologies, LLC. Los Angeles, California). Each participant’s raw data were digitally band-pass filtered by a linear-phase FIR filter (0.01-0.1 Hz) to filter noises due to heart beat (0.8~1.2 Hz), respiration (0.3 Hz) and Mayor waves (≥ 0.1 Hz) (32, 33). Then, each wavelength was detrended by their respective whole length record (without segmentation), and variations in concentration of oxyhemoglobin and deoxyhemoglobin were calculated by the modified Beer-Lambert law (differential pathlength factor set to 7.25 and 6.38, respectively) (34). Each concentration curve was then segmented into the 30 s of interest of each trial, for all studied conditions.

The mean concentration of oxyhemoglobin and deoxyhemoglobin for each segment was calculated for each channel using the average of moving 2s-window means with 50% overlap. Thus, each subject’s database was composed of 64 features of average concentration (32 channels × 2 chromophores) in 20 experimental conditions (10 neutral trials + 5 positive trials + 5 negative trials), for both tasks.

#### 2.4.2. Machine learning procedure

To decode affective states across databases, we used the Linear Discriminant Analysis (LDA) implementation provided by the BCILAB toolbox (35). To avoid overfitting the model to our dataset, the LDA was computed with its default parameters. This analysis was divided in two main steps represented in Figure 3.

**Figure 3.**
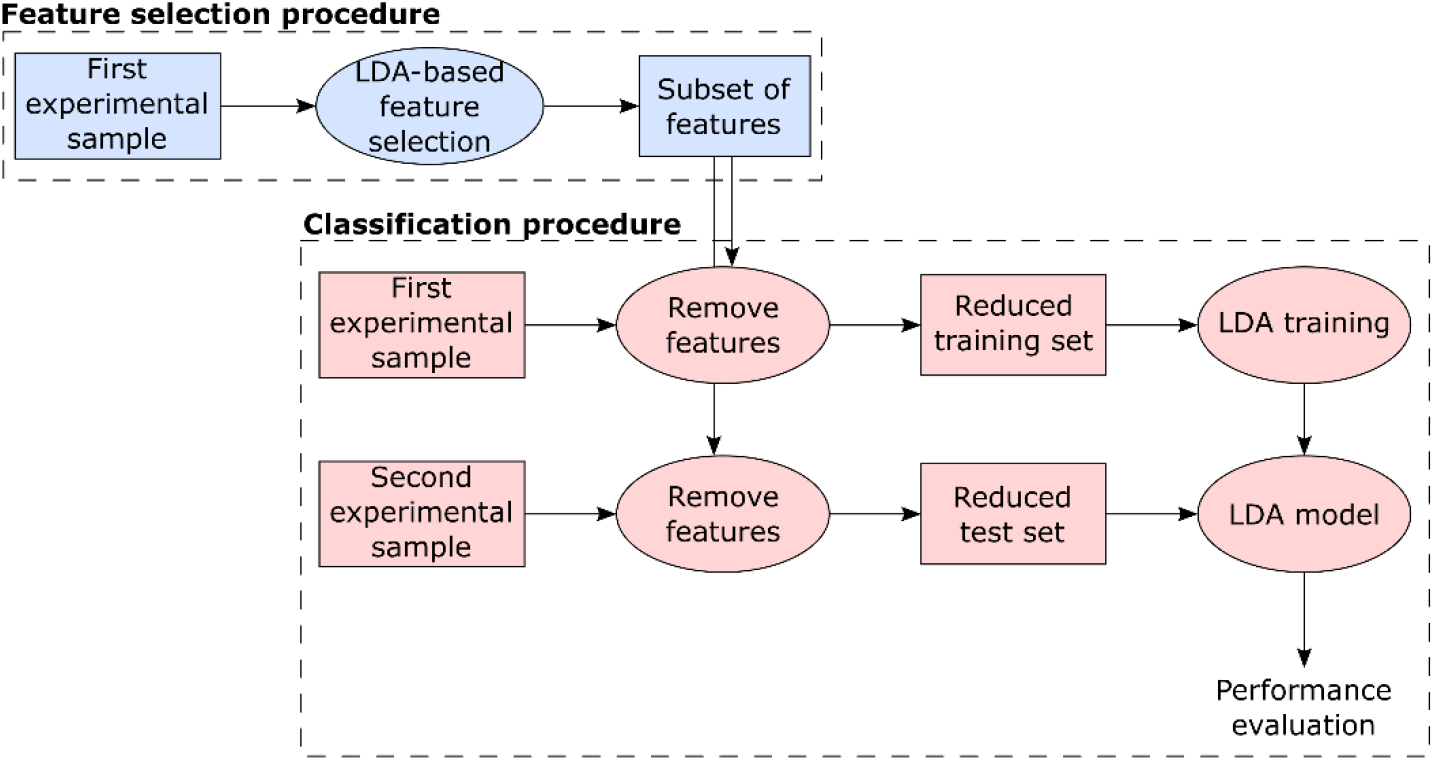
Machine learning procedure: Initially, we performed an LDA-based feature selection using the first experimental sample (49 subjects). Then, based on these selected features, we created training and test sets for the LDA classifier using the first (49 subjects) and second (20 subjects) experimental samples, respectively.

First, we performed an LDA-based feature selection using the whole feature space from the first experimental sample (49 participants) (steps in blue in Figure 3). The LDA model searches for a linear projection of multivariate observations to univariate ones (36). Thus, it is possible to use the eigenvalues of the within-class covariance matrix in LDA (which contain most of the relevant information) as a measure of feature relevance (37). To reduce the number of features used during classification, we first ranked the absolute eigenvalues from all features, and then created different subsets selecting the best 5 to 100% (with steps of 5%) features with highest eigenvalues (in other words, one subset using the top 5% features, another subset using the top 10%, and so on).

For each obtained feature subset, projections of the training and testing datasets described by the selected features only were built using the first (49 participants) and second (20 participants) experimental samples, respectively (steps in red in Figure 3). Then, we evaluated the predictive performance on each corresponding test set. This approach was applied to evaluate affective conditions in pairs, following “Positive versus Neutral”, “Negative vs. Neutral” and “Positive vs. Negative” combinations, for reactive and active tasks separately. Here we followed this binary approach since many current neurofeedback protocols are mainly based on two-class designs (24, 38–43).

To evaluate the prediction performance in each comparison, we calculated the decoding accuracy as (trials correctly classified from first class + trials correctly classified from second class)/2. This approach avoids the potential unbalance of sample size in each comparison, and this random result should be 50%.

#### 2.4.3. Statistical analysis

Considering that the results here presented are sum-based proportions, and no ceiling-effect was observed, these data assume approximately normal distributions. Thus, here we applied parametric t-test to evaluate our results.

First, the subject scores of valence and arousal of the second database were evaluated by a two-sample t-test. The comparisons used the mean values of valence and arousal assigned during positive, negative and neutral trials. This procedure was repeated for both tasks. The p-values were Bonferroni corrected for multiple comparisons (2 subjective measures × 3 comparisons of conditions × 2 tasks).

The significance of classification accuracy of each comparison was evaluated by a one sample t-test, comparing the prediction performance from all participants in this comparison against the chance level (50%). Each result was independently adjusted by using the Bonferroni correction for 120 multiple comparisons (20 different percentages of features × 3 comparisons of conditions × 2 tasks = 120).

## 3. Results

### 3.1. Subjective scores of Arousal and Valence

Table 1 presents the mean subjective scores of arousal and valence relative to the positive, negative and neutral conditions, during both active and reactive tasks. Moreover, the corrected p-values of each comparison are also presented.

**Table 1.** In the first half of the table, mean ± standard-deviation of subjective scores (ranging from 1 to 9) assigned by each subject of the second database during negative, neutral and positive trials. In the second half, p-values of the comparison between the scores assigned for each trial type.

| Mean subjective scores |  |  |
| --- | --- | --- |
|  | Valence | Arousal |
| **Reactive task** |  |  |
| Negative | 2.52 ± 1.38 | 6.05 ± 1.34 |
| Neutral | 5.11 ± 0.73 | 2.71 ± 1.29 |
| Positive | 6.79 ± 0.92 | 4.82 ± 1.60 |
| **Active task** |  |  |
| Negative | 2.57 ± 0.97 | 5.85 ± 1.71 |
| Neutral | 4.84 ± 0.24 | 2.09 ± 1.39 |
| Positive | 7.20 ± 0.77 | 5.67 ± 1.75 |

| Statistical comparisons (p-values) |  |  |
| --- | --- | --- |
|  | *Valence* | *Arousal* |
| ***Reactive task*** |  |  |
| Positive vs. Negative | <0.01 | 0.14 |
| Positive vs. Neutral | 0.02 | 1.00 |
| Negative vs. Neutral | <0.01 | <0.01 |
| ***Active task*** |  |  |
| Positive vs. Negative | <0.01 | 1.00 |
| Positive vs. Neutral | <0.01 | <0.01 |
| Negative vs. Neutral | <0.01 | <0.01 |

Valence scores presented significant differences for all the comparisons. It is relevant, however, that no significant difference of the arousal scores was found during “Positive vs. Negative” comparisons, as well as during “Positive vs. Neutral” comparison during the reactive task.

### 3.2. Classification results

Boxplots describing the LDA accuracy are presented in Figure 4. Classification accuracy for the reactive task significantly exceeded chance level in “Positive vs. Negative” comparisons, with highest result as 64.50 ± 12.03% (mean ± standard-deviation, p<0.01) using 20% of features (12 channels), and in “Negative vs. Neutral” comparisons (68.25 ± 12.97%, p<0.01, 30% of features – 19 channels). No significant differences were found for “Positive vs. Neutral” comparisons.

**Figure 4.**
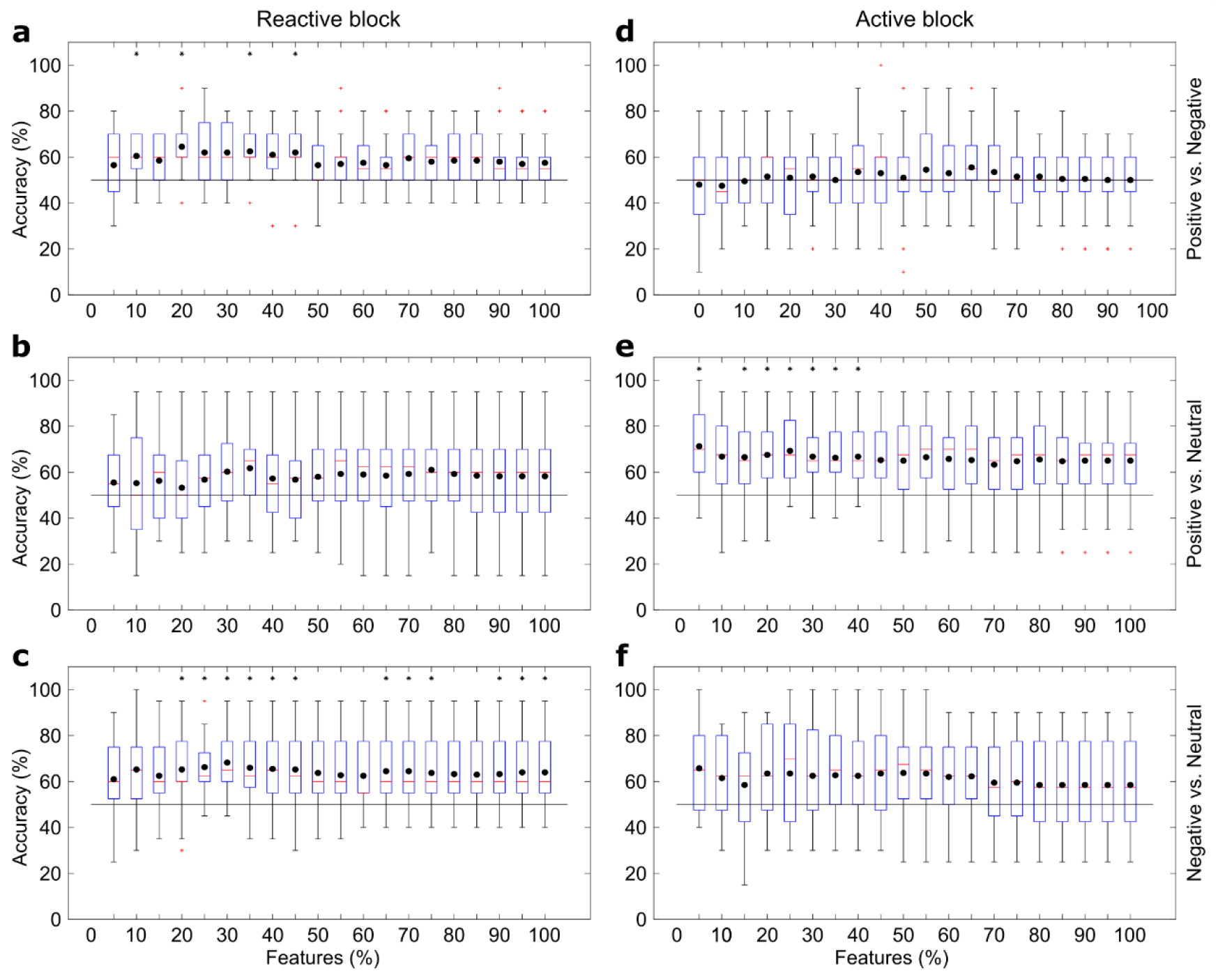
Box plots showing the results using the LDA classifier and different feature subsets. (a) presents the “Positive vs. Negative”, (b) the “Positive vs. Neutral”, and (c) the “Negative vs. Neutral” comparisons for reactive elicitation block, while (d-f) follows the same order for active elicitation block.Black dots present the means, red lines the medians, red crosses the outliers and black asterisks the statistical difference for chance level (p<0.05).

For the active task, accuracies were greater than chance level in “Positive vs. Neutral” comparisons, and the highest result was 71.25 ± 18.02% (p<0.01) using only 5% of the features (3 channels). No significant differences were found for “Positive vs. Negative” and “Negative vs. Neutral” comparisons.

### 3.3. Relevant channels

In Figure 5, the channels selected in each of the highest significant results reported in Section 3.2 are shown alongside their respective weights assigned by the LDA classifier. The location of the channels follows the same order shown in Figure 1.

**Figure 5.**
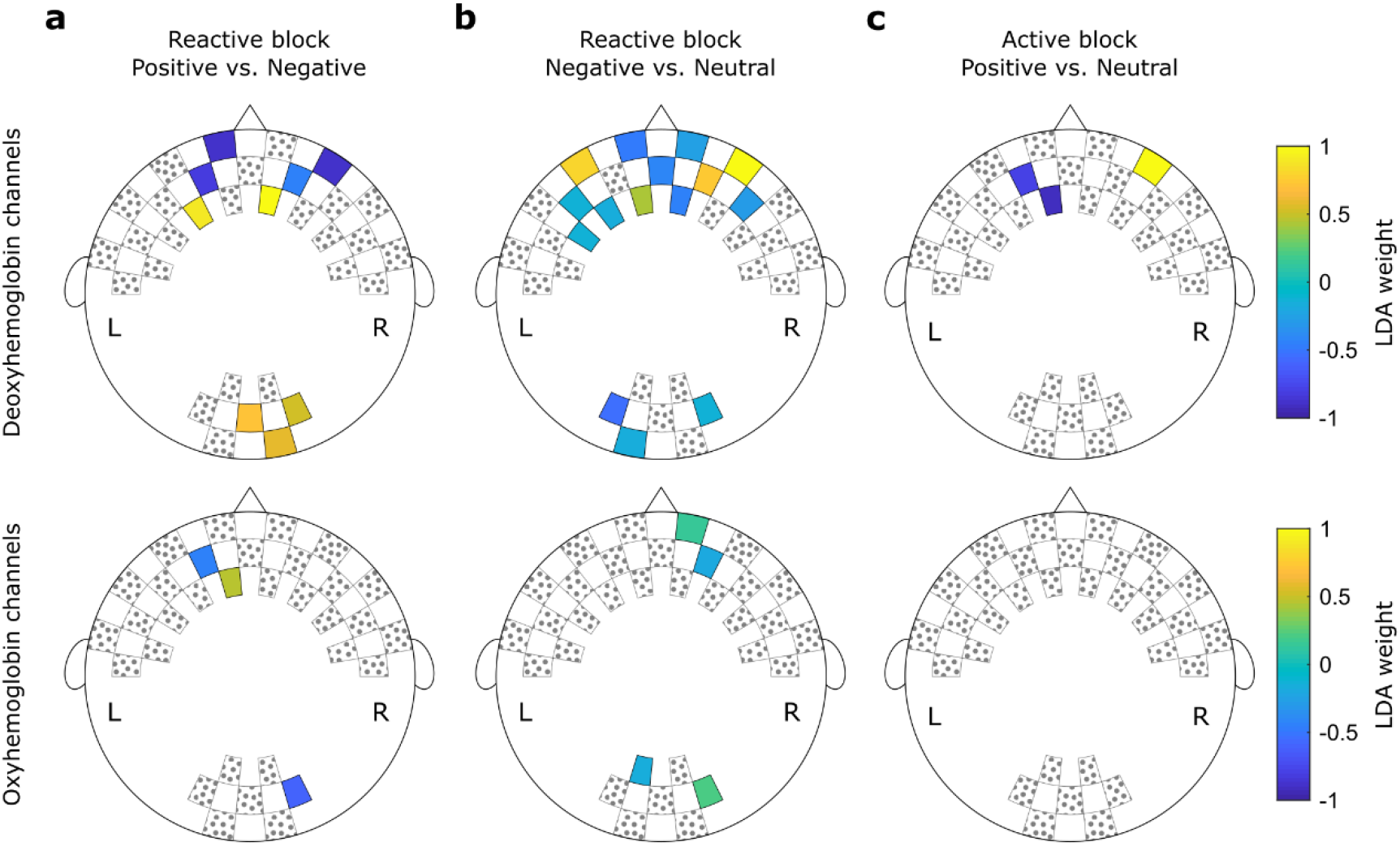
Weights assigned by the LDA classifier for each feature during the (a) “Positive vs. Negative” comparisons using 20% of features and (b) “Negative vs. Neutral” comparisons using 40% of features. In (c), the weights for each feature during the “Positive vs. Neutral” comparisons using 5% of features. Hottest colors indicate positive weights while cooler colors indicate negative weights. Channels filled with gray dots were not used during the test.

Among the “Positive vs. Negative” classification during the reactive task, the highest accuracy was achieved using 12 features. Nine of these features correspond to information about the deoxyhemoglobin concentration. According to the 10-5 EEG electrode positioning system (44), four channels are approximately at the frontopolar area, two at the anterofrontal area, and three at the occipital area. Complementary, three features included information about the oxyhemoglobin concentration at the frontopolar (two channels) and the occipital (one) areas.

For the “Negative vs. Neutral” classification during the reactive task, 19 channels carried relevant information: six frontopolar, five anterofrontal, one frontal, and three occipital channels for the deoxyhemoglobin concentration; and two frontopolar and two occipital channels for the oxyhemoglobin concentration.

Finally, the “Positive vs. Neutral” decoding during the active task used only three channels to achieve the highest performance: two frontopolar and one anterofrontal. These channels provided information about the deoxyhemoglobin concentration. The corresponding averaged time series are shown in Figure 6, as well as the averaged values used as inputs to the LDA classifier. It is notable that the time series from second sample have higher variability, since the number of subjects (20) is smaller than the first sample (49). However, it is also notable that the averaged features follow the same patterns in both samples, with positive trials presenting higher concentration levels in channel 6-7, and neutral trials presenting higher concentrations in channels 5-4 and 4-4. This is also consistent with the directions of the LDA weights showed in Figure 5c.

**Figure 6.**
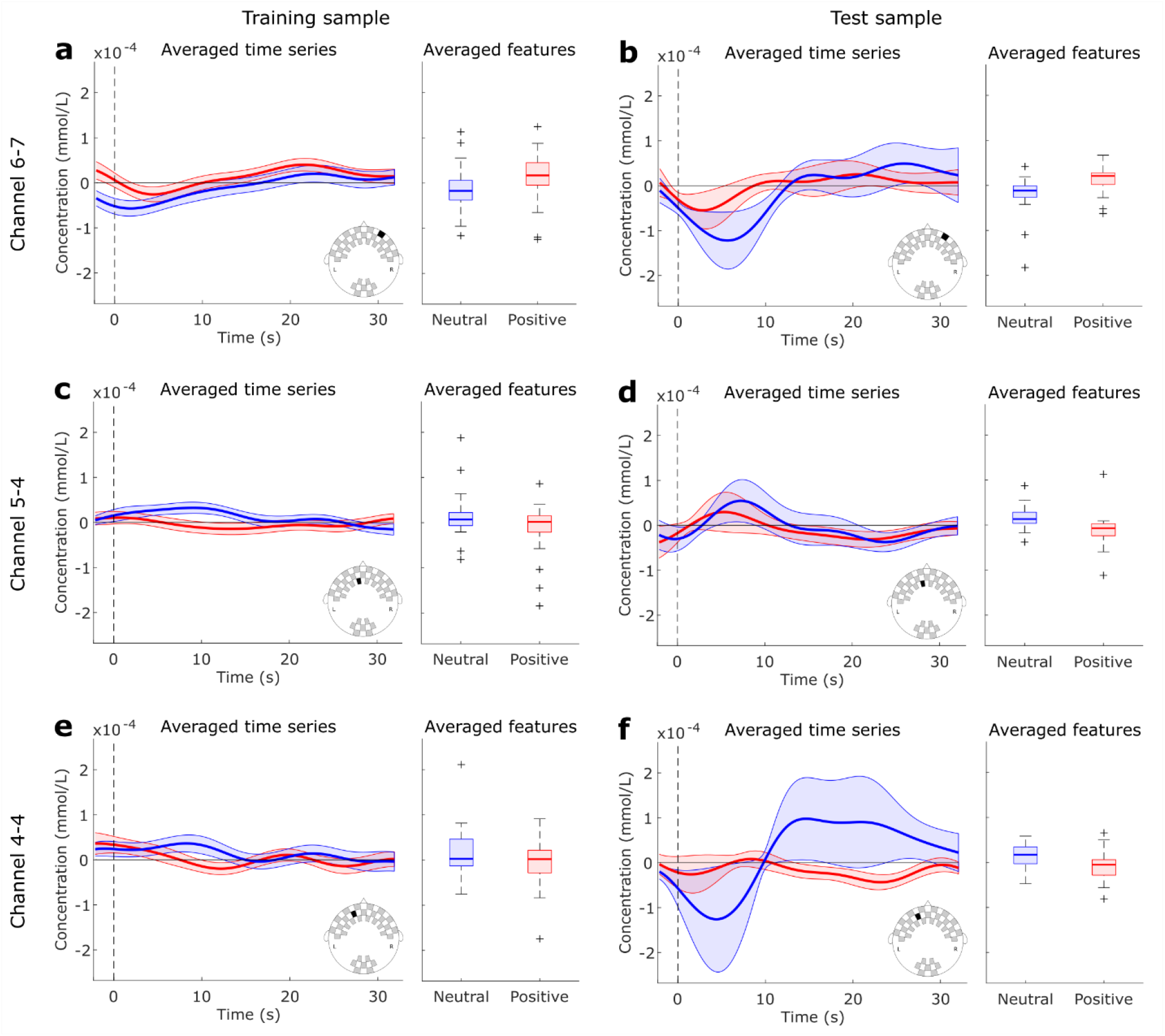
Averaged time series and features used as inputs to the LDA classifier during “Positive vs. Neutral” comparison in the active task. Data from Channel 6-7 is shown in (a-b), Channel 5-4 in (c-d) and Channel 4-4 in (e-f) for the training and test samples, respectively. Blue curves and boxes represent Neutral trials, and red curves and boxes Positive trials.

## 4. Discussion

### 4.1. Affective decoding

First, it is essential to highlight the absence of statistical significance in the subjective arousal score during the “Positive vs. Negative” comparisons. In addition to the significant difference in the valence scores, it suggests that any classification result in these comparisons is exclusively related to valence differences between positive and negative affect. Further, during both “Positive vs. Neutral” and “Negative vs. Neutral” comparisons, statistical differences were found for valence and arousal scores. It is expected that Positive and Negative present differences in arousal and valence when compared to absent affective content (Neutral). Some studies suggest valence and arousal not as two orthogonal dimensions, but as a single V-shaped dimension (45). Accordingly, these two dimensions cannot be separated in most of the emotional stimuli systems (for example, the International Affective Pictures System) (27, 46). Given this, we have demonstrated that it is possible to detect distinct patterns of hemodynamic activity generated by different affective valence elicitation tasks. Our results also provide evidences for the feasibility of decoding affective states in a new, untested subject, based on a model trained in previously collected and independent dataset.

For the decoding of the reactive task, only the accuracies for “Positive vs. Negative” and “Negative vs. Neutral” comparisons were higher than chance level. This finding is in line with previous inter-subject studies using the IAPS database to decode affective states, but with fMRI (13). In a qualitative evaluation, some participants described, after the experiment, that the negative affect induced by the pictures presented during the reactive block was more intense than the positive one. Negative figures into the IAPS catalog are mainly related to death, malnutrition, sickness, poverty, and disgust (25), which are more consensual than the content of some positive stimuli, such as babies, pets or beaches. Moreover, some studies suggest that processing negative stimuli are more demanding in the brain (47), which might generate more clear signals to our classifiers than the neural processing of positive stimuli.

The decoding of the active task, on the other hand, presented significant accuracy for the “Positive vs. Neutral” condition exclusively. The participants also described that the positive imagery during active condition was more natural to achieve and more intense than the negative representation. This result is particularly interesting for potential neurofeedback applications (4, 48). For example, in psychiatric applications where patients present increased levels of negative emotions and decreased levels of positive emotions (49), improving the self-regulation between Positive and Neutral states might be clinically relevant (24).

### 4.2. Relevant features and the neural networks of affect

For both results in the reactive task, relevant features included both fNIRS derived oxyhemoglobin and deoxyhemoglobin concentrations measures. In fact, each chromophore is expected to provide different and complimentary information. Deoxyhemoglobin is thought to be an indicator of functional activation and is more directly related to the fMRI BOLD signal (15, 50), while a recent work found a correlation between oxyhemoglobin signal and the EEG band power variation in some cortical regions (51). Other decoding experiments also reported both hemoglobin concentrations as relevant for classification (19). These results suggest a non-redundancy of these measures.

In accordance with two meta-analyses of neuroimaging data in affective tasks, we found that occipital areas signals were among the relevant features for classification (28, 29). Additionally, fNIRS studies also described activation of the occipital cortex during affective stimulation (52, 53). Minati and colleagues, for example, found increased occipital response to positive and negative affects relative to neutral pictures of the same database we have used (IAPS) (54). Moreover, it is interesting to note that the classifier assigned high absolute weights for a considerable number of channels from frontopolar and anterofrontal regions. A frontal lateralization of activity during the experience and regulation of emotional responses is a classical effect (55–58). However, a more broad theoretical account of affective processing, affective workspace hypothesis, states that activity patterns of the same core neural network could implement both positive and negative affective responses (26, 27). Our application of subject-independent affective decoding, which considers distributed brain activity, is in line with the theoretical assumptions of such model. Frontal regions, which were consistently relevant for classification in our experiments, superpose to the network nodes that have been proposed as core regions of the affective neural workspace (28, 59).

### 4.3. Subject-independent designs

fNIRS-based affective neurofeedback protocols were recently applied to healthy (24, 60) and psychiatric populations (61), including patients with schizophrenia (62), autism disorder (63), and attention-deficit/hyperactivity disorder (ADHD) (64, 65). However, all these protocols have subject-specific designs which require training blocks or calibration trials for every experimental session. In addition to the expected duration of the setup preparation, these demands can act as stressors for patients presenting anxiety symptoms, or cause physical and mental fatigue shortly after the initial blocks (6). Consequently, patients may not achieve their best performance, or the therapeutic benefits associated with the protocol. All these aspects make the investigation of subject-independent fNIRS protocols a timely research topic.

Critically, classification accuracies presented in this paper are lower than those presented in our previous subject-specific study (22). However, the performance drop during subject-independent designs is an expected effect due to the inter-subject variability of anatomy, level of stress, rest, among others. For example, Robinson and colleagues report more than 10% of performance reduction from a subject-dependent to a subject-independent fNIRS-based motor imagery neurofeedback (66).

Also, the mean accuracy for “Positive vs. Neutral” discrimination in the active block was slightly over the 70% threshold suggested by the brain-computer interface and neurofeedback communities as sufficient to perform device control and communication (19, 67). This is a crucial finding since the differentiation between the active elicitation of positive affect and a neutral (resting-state) condition is the state of the art of hemodynamic-based neurofeedback protocols applied to both health and psychiatric populations (24, 39, 41, 42, 68, 69). It is also important to emphasize that this result was reached using only three channels. Considering future applications, the use of only three channels means shorter setup procedure, lower instrumental and computational costs and the possibility of even more portable systems.

### 4.4. Limitations and future perspectives

Despite the controls we implemented in our experimental design, collection and analysis, limitations should be recognized. First, although fNIRS signal is mainly related to the near-infrared light absorption into the cortical surface, it also encompasses peripheral responses, including changes in skin, muscular and cranial blood flow and the aerobic process of energy consumption related to muscle contractions (16). Consequently, some authors refer to fNIRS applications for control of devices such as corporeal machine interfaces (19). As this constitutes one fundamental issue of this method, further validation using this approach is warranted. Here, one approach that could be explored in future studies is the use of short-separation channels to filter systemic hemodynamic fluctuations from non-neural sources (70, 71).

Also, we used minimal preprocessing steps, classifying fNIRS data using the mean changes in oxy and deoxyhemoglobin concentrations as features. This approach was used in order to test fNIRS based MDB of affective state with the least assumptions possible, to avoid introducing spurious artifact in the signal and to test the feasibility for posterior real-time analysis (72, 73). In comparison, recent studies identified that combining the mean hemoglobin concentration with other temporal and time-frequency features improves the decoding accuracies reaching values close to 90% in within-subject decoding (19, 74, 75). Therefore, future studies should also evaluate the effect of different feature extraction techniques to the inter-participants MBD of affective states.

However, even with these limitations, we reinforce the advantages of fNIRS compared to other neuroimaging techniques in implementing MBD systems. Particularly, the portability, benefit-cost relation and the good balance between spatial and temporal resolutions makes fNIRS especially suitable for this purpose (16). In fact, these aspects create a solid base for an increasing number of studies using fNIRS for MBD of motor, cognitive, and more recently for decoding of affective states (19, 20, 22). In subject-independent designs, experiments can also be found classifying semantic experiences (76) or audiovisual processing (77). Additionally, the portability of fNIRS systems allow a fast evolution of naturalistic experiments targeting real-world applications (78, 79), such as affective neurofeedback systems for therapeutic purposes (24).

## 5. Conclusion

Although more experiments are necessary to increase the classification accuracies reached here, our results suggest that fNIRS is a potential tool for subject-independent decoding of affective states. The accuracy in discriminating positive and neutral emotions during the affective task was significant and above the threshold desired for effective control of BCI or neurofeedback protocols. Thus, the construction of fNIRS-based subject-independent neurofeedback devices should be attempted.

## Acknowledgements

The authors thank Guilherme A. Z. Moraes (NIRx) and Jackson Cionek (Brain Support Brazil) for the technological support. This study was funded by the São Paulo Research Foundation (FAPESP): LRT received grant number 2015/17406-5, JT received grant number 2017/05225-1, JRS received grants number 2018/21934-5 and 2018/04654-9. The funders had no role in study design, data collection and analysis, decision to publish, or preparation of the manuscript.

The authors declare no conflict of interest.

